# Imitation as a model-free process in human reinforcement learning

**DOI:** 10.1101/797407

**Authors:** Anis Najar, Emmanuelle Bonnet, Bahador Bahrami, Stefano Palminteri

## Abstract

While there is not doubt that social signals affect human reinforcement learning, there is still no consensus about their exact computational implementation. To address this issue, we compared three hypotheses about the algorithmic implementation of imitation in human reinforcement learning. A first hypothesis, *decision biasing*, postulates that imitation consists in transiently biasing the learner’s action selection without affecting her value function. According to the second hypothesis, *model-based imitation*, the learner infers the demonstrator’s value function through inverse reinforcement learning and uses it for action selection. Finally, according to the third hypothesis, *value shaping*, demonstrator’s actions directly affect the learner’s value function. We tested these three psychologically plausible hypotheses in two separate experiments (N = 24 and N = 44) featuring a new variant of a social reinforcement learning task, where we manipulated the quantity and the quality of the demonstrator’s choices. We show through model comparison that *value shaping* is favored, which provides a new perspective on how imitation is integrated into human reinforcement learning.

## Introduction

The complexity of our society could not have been achieved if humans had to rely only on individual learning to identify solutions for everyday decision problems. An isolated learner must invest sufficient energy and time to explore the available options and may thus encounter unexpected negative outcomes, making individual learning costly, slow and risky. Social learning may largely mitigate the costs and risks of individual learning by capitalizing on the knowledge and experience of other subjects.

Imitation is one main mechanism of social learning and has been widely investigated in psychology and neuroscience (1, 2). It has been studied from various frameworks, such as the mirror neurons system (3), theory of mind (4) and Bayesian inference (5). While these studies provide valuable insights about the computational mechanisms of imitation, they are still limited in their scope as imitation is treated in isolation from other learning processes, such as autonomous - reinforcement - learning. Here we compare three radically different and psychologically plausible computational implementations of how imitation can be integrated into a standard reinforcement learning algorithm (6–9). To illustrate the three hypotheses we consider the stylized situation, where a reinforcement *Leaner* is exposed to the choices of a *Demonstrator*, before making her own choices.

The first hypothesis, *decision biasing*, well represented in the cognitive neuroscience literature, postulates that observing a *Demonstrator*’s action influences the learning process by biasing the *Learner*’s decision on the next trial towards the observed action (6, 7). This implementation presents one limitation in that it does not allow an extended effect of imitation over time. This hypothesis conceptualizes imitation as biasing the exploration strategy of the *Learner* and contrasts with the repeated observations in experimental psychology suggesting that social signals have long-last effects on the *Learner*’s behavior (10). In addition, the experimental designs of these previous studies were not well suited for assessing whether imitation integrates over successive demonstrations and propagates over several trials, since one observational trial was strictly followed by one private trial.

A second account consists in framing imitation as an inverse reinforcement learning problem, where the *Learner* infers the preferences of the *Demonstrator* (5, 9). In previous paradigms, the model of the *Demonstrator* is generally used to predict her behavior, but these representations could easily be recycled to influence the behavior of the *Learner*. Unlike the *decision biasing* account, this *model-based imitation* allows for the accumulation of demonstrations and the propagation of imitation over several trials. However, this method is computationally more demanding and still leaves unanswered the question of how the model of the *Demonstrator* is integrated into the behavior of the *Learner*.

In line with previous works on advice taking (11) and learning from evaluative feedback (12), we propose a third approach where the demonstrations influence the value function instead of action selection. As such, this *value shaping* scheme has the desirable properties of allowing for a long lasting influence of social signals while being computationally simple.

We compared these three computational implementations of imitation (and a baseline model without social learning; see Fig. 2) against empirical data from two independent experiments (Exp.1: N = 24, Exp.2: N = 44). Both experiments featured a new variant of a social reinforcement learning task, where, to assess the accumulation of social signals, we manipulated the number of demonstrations in a trial-by-trial basis (Fig. 1). We also manipulated the skills of the *Demonstrator* to assess whether the imitation parameters were modulated in an adaptive manner. This manipulation was implemented within-subject in Exp.1 and between-subject in Exp.2, to see whether observing both *Skilled* and *Unskilled Demonstrators* had a different effect on performance than observing only a *Skilled* or an *Unskilled Demonstrator*.

**Fig. 1.**
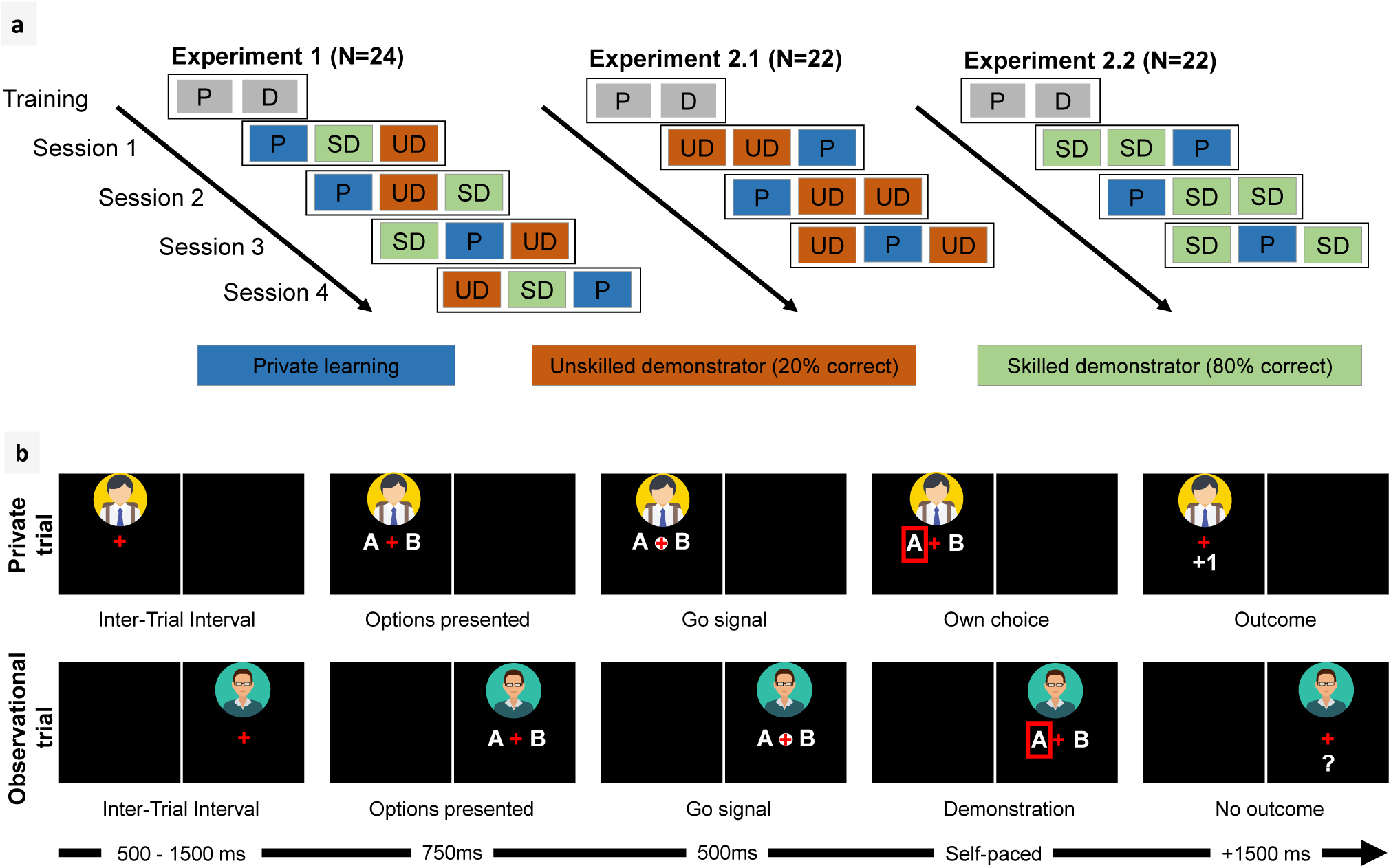
Experimental design. (A) Experimental conditions. Conditions were randomized within sessions. In the Private condition (P), there was no social information and the task was a basic reinforcement learning task. In observational conditions (D) the *Learner* observed choices of another player that were inter-leaved with her own choices. The training involved a short Private session followed by an Observational session. In the *Skilled Demonstrator* condition (SD), the other player picked the reward maximizing symbol 80% of the time, while in the *Unskilled Demonstrator* condition (UD), the other player picked the reward maximizing symbol only 20% of the time. The correct choice rate of the other player (SD-UD contrast) was manipulated within-subject in Exp.1, and between-subjects in Exp.2. Experiments 1 and 2 comprised 4 and 3 sessions respectively. (B) Typical observational and private trials (timing of the screen given in milliseconds). During observational trials, the *Learner* was asked to match the choice of the other player before moving to the next trial.

**Fig. 2.**
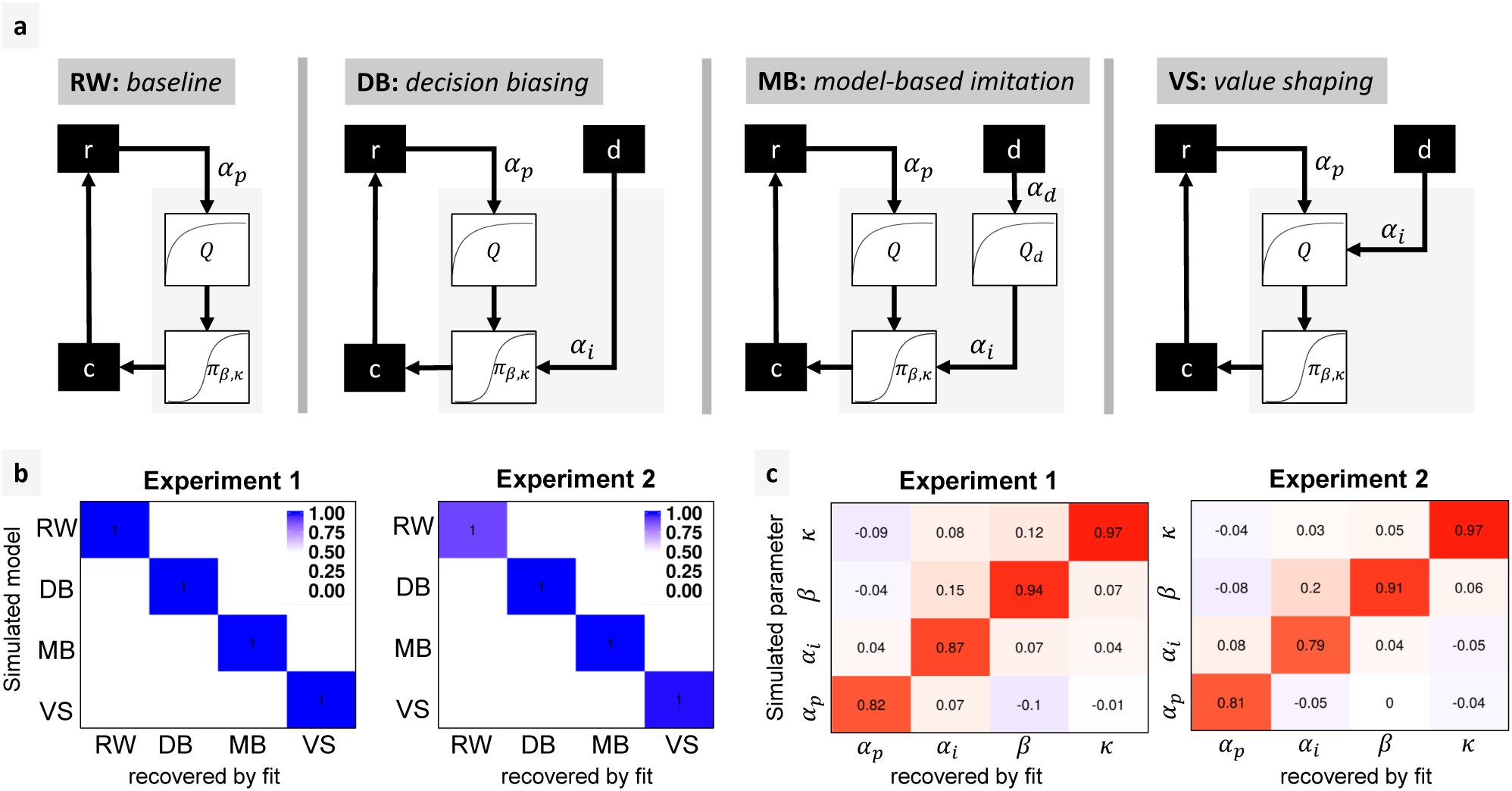
Model comparison. a) Model space: RW: the baseline is a Rescorla-Wagner model that only learns from self-experienced rewards. Rewards *r* are used for computing a value function *Q*, which in turn is used for deriving a policy *π* using a soft-max function. Subject choices *c* are sampled according to *π*. The demonstrations *d* have no effect on the learning process. DB: *decision biasing*. Demonstrations are used for biasing the subject’s policy without affecting the value function. MB: *model-based imitation*. Demonstrations are used for inferring the *Demonstrator*’s value function *Q*_*d*_, which is then used for biasing the subject’s decisions. VS: *value shaping*. Demonstrations directly affect the subject’s value function. *α*_*p*_ private reward learning rate. *α*_*i*_ imitation learning rate. *α*_*d*_ demonstration learning rate. *β* soft-max inverse temperature. *κ* choice auto-correlation. b) Model recovery: model frequencies (blue shade) and exceedance probabilities (XP = 1) for each pair of simulated/fitted model, based on the AIC. c) Parameter recovery: Spearman correlation scores between each pair of simulated/fitted parameter values for the winning model VS.

Overall, quantitative model comparison indicated that imitation, whenever adaptive (*i.e.*, when the *Demonstrator* outperforms the *Learner*), takes the computational form of *value shaping*, rather than *decision biasing* or *model-based imitation*.

We also analyzed the parameters of the best winning model to determine whether or not, in the context of reinforcement learning, imitation is subject to an *egocentric* bias, where more weight is given to information derived from oneself compared to the other (13). The comparison of the private reward learning rate with the imitation learning rate was overall consistent with an *egocentric* bias. Finally, we compared the imitation learning rates across different observational condition (*Skilled* vs. *Unskilled Demonstrators*), to see whether imitation is modulated. Consistent with our model comparison results and with previous theoretical and empirical work about meta-learning (8, 14), we found that subjects can infer the skill of a *Demonstrator* and regulate imitation accordingly. This result was even more pronounced when the skill of the *Demonstrator* was implemented within-subject, which suggests that being exposed to both *Skilled* and *Unskilled Demonstrators* models could be more effective in preventing *Learners* from bad influence.

## Results

### Experimental design

We performed two experiments implementing a probabilistic instrumental learning task (cf. Fig. 1), where the participants were repeatedly presented with a binary choice between two abstract visual stimuli resulting in either winning or losing a point. Displayed stimuli had opposite winning and losing probabilities (0.7/0.3 in Exp.1 and 0.6/0.4 in Exp.2). The goal of the participant, *i.e.*, the *Learner*, was to learn by trial-and-error which of the two stimuli had the highest expected value. Three learning conditions were presented in blocs of 20 trials each, per session. In the *Private* condition (P), no information other than reward outcome was provided to the *Learner*. Observational conditions involved additional social information that was randomly interleaved within private choices. Specifically, during observational trials the *Learner* could observe the choice of another player, hereafter referred to as the *Demonstrator*, on the same pair of stimuli. Participants were recruited in pairs, and told that they were observing the choices of the other participant. However, the demonstrations were controlled by the experimenter and drawn from two Bernoulli distributions. In the *Skilled Demonstrator* condition (SD), the other player picked the reward maximizing stimulus 80% of time. In the *Unskilled Demonstrator* condition (UD), the other player picked the reward maximizing stimulus only 20% of time. The SD/UD manipulation was within-subject in Exp.1 (all subjects were exposed to both *Skilled* and *Unskilled Demonstrators*) and between-subject in Exp.2 (each subject was exposed to either the *Skilled* or the *Unskilled Demonstrator*).

### Behavioral analyses

As a quality check, we first assessed whether or not subjects overall learned to choose the correct option. We found the average correct choice rate significantly above chance level in both experiments (Exp.1: M = 0.75, SD = 0.08, t(23) = 14.85, p = 2.8e-13; Exp.2: M = 0.62, SD = 0.09, t(43) = 8.56, p = 7.69e-11), indicating that subjects overall identified the reward maximizing symbols. We then looked at whether or not the correct choice rate was affected by the skill of the *Demonstrator*. In Exp.1, even if as expected the average correct choice rate was higher in the SD compared to the UD condition (76% vs. 73%), the difference did not reach statistical significance (*F* (2, 69) = 0.426, *p* = 0.655). In Exp.2, we found a significant interaction between *Demonstrator*’s performance and social information (*F* (1, 84) = 4.544, *p* = 0.036). The effect was driven by *Learners* in the SD group having a correct choice rate differential greater than zero (+5%) comparing the Observational and Private conditions, and *Learners* in the the UD having a differential smaller than zero (−4%) (direct comparison of the learning differential between UD vs. SD: Welch Two Sample t-test *t*(35.215) = 2.8589, *p* = 0.007, cf. supplementary Fig. 1). These results indicate that *Demonstrators*’ skill affected *Learners*’ performance.

### Model comparison

We fitted four computational models to the behavioral data of both experiments (Fig. 2a). In our model space, the baseline was represented by a *Rescorla-Wagner* model (RW) that does not integrate social information into the learning process (*i.e*, it treats Private and Observational conditions equivalently). The second model, *decision biasing* (DB), assumes that demonstrations only bias action selection on the next trial (6). The third model, *modelbased imitation* (MB), implements an inverse reinforcement learning process that infers a model of the *Demonstrator*’s preferences and uses it for biasing action selection. Finally, in the *value shaping* model (VS), *Demonstrator*’s choices directly affect the value function of the *Learner*. The exact computational implementation of each model was first optimized independently, by comparing several alternative implementations of the same model: two implementations of RW, six for DB, nine for MB and two for VS (cf. Methods). In the main text, we only report the comparison between the best implementations of each model. The intermediate model comparison results are provided with the supplementary materials (cf. supplementary Fig. 2).

We first performed a model recovery analysis to check the models are identifiable within our task and parameter space (Fig. 2b). We then compared the different models based on the experimental data (Fig. 3a). In Exp.1, roughly the same proportion of subjects were explained by RW (Model Frequency MF: 0.45, Exceedance Probability XP: 0.72) and VS (MF: 0.33, XP: 0.24). These results can easily be explained by the fact that all participants encountered both *Skilled* and *Unskilled Demonstrators*. Since *Unskilled Demonstrators* should not be imitated, their presence can justify the observed high frequency of RW. We verified this intuition by fitting separately UD and SD conditions (plus the Private condition). Indeed, we observed that subjects’ behavior in the UD condition was best explained by RW (MF: 1, XP: 1), whereas the choice data in the SD condition were better explained by VS (MF: 0.9, XP: 1).

**Fig. 3.**
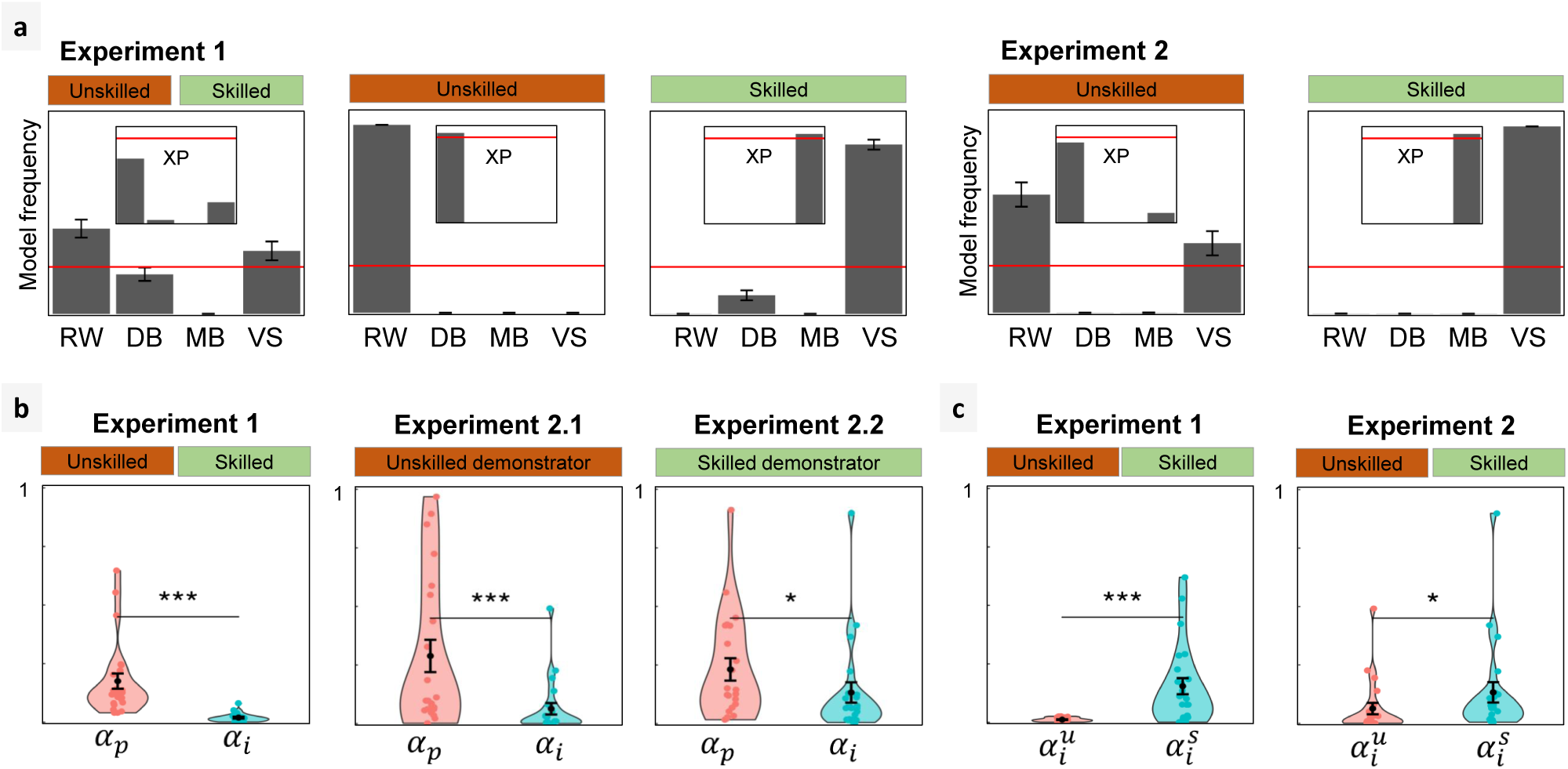
a) Model comparison: model frequencies and exceedance probabilities (XP) of the fitted models based on the AIC. Red lines: 0.25 chance level for model frequency and 0.95 threshold for exceedance probability. b,c) Learning rates of the winning model VS. b) comparison between private (*α*_*p*_) and imitation (*α*_*i*_) learning rates. b) comparison between imitation learning rates when observing the *Unskilled* 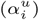 and the *Skilled* 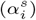 *Demonstrator*.

The same pattern was found in Exp.2 where the skill of the *Demonstrator* was manipulated between subjects. The subgroup of subjects in the UD condition were best explained by the RW model (MF: 0.63, XP: 0.89), while the subgroup of subjects in the SD condition were best explained by the VS model (MF: 1, XP: 1).

Of note, the model comparison results were robust across different implementations of the model space, notably with or without asymmetric value update in private learning, with or without including a choice auto-correlation parameter, and with or without allowing for negative imitation learning rates (cf. Methods and supplementary Fig. 3).

To sum up, in both experiments we consistently found that, when useful, imitation is computationally implemented in a *value shaping* way. These results show that in our task, imitation has a direct effect on the value function, by reinforcing the observed actions, but without building an explicit model of the *Demonstrator*.

### Parameter analysis

After verifying that the parameters of the winning model, VS, can be properly identified (Fig. 2c), we analyzed the fitted parameters across experiments and conditions (Fig. 3b,c).

We first assessed whether subjects overweighted their freely obtained outcomes (*egocentric bias*) or the choices of the *Demonstrator* (*allocentric bias*). Consistent with the *egocentric bias* hypothesis, we found the private reward learning rates significantly higher compared to the imitation learning rates (Fig. 3b). This was true in Exp.1 (Wilcoxon signed-rank test: *V* = 300, *p* = 1.192*e*−07). In Exp.2, the difference between private and imitation learning rates was even more pronounced when confronted with an *Unskilled Demonstrator* (Wilcoxon signed-rank test: *V* = 251, *p* = 1.431*e*−06), and still detectable when facing a *Skilled Demonstrator* (Wilcoxon signed-rank test: *V* = 200, *p* = 0.01558).

We then compared imitation learning rates across different observational conditions to see whether imitation was modulated by the skill of the *Demonstrator* (Fig. 3c). We found the imitation learning rate significantly higher in the SD condition compared to the UD condition (Exp.1: Wilcoxon signed-rank test: *V* = 9, *p* = 3.934*e* − 06, Exp.2: Wilcoxon rank-sum test: *W* = 137, *p* = 0.01312). We also inspected the other (non-social) parameters such as choice temperature and private reward learning rate to see whether the effect of the *Demonstrator* skill was specific to imitation. This analysis revealed no significant difference between the non-social parameters fitted in the UD and the SD conditions (Exp.1: all *p >* 0.09; Exp.2: all *p >* 0.2; cf. supplementary Fig. 4). This suggests that the strategic adjustment induced by the *Demonstrator*’s skill is specific to the imitation process. The result of this analysis is consistent with the model comparison results indicating that different computational models (RW and VS) explain the *Learner*’s behaviour in the different observational conditions (UD and SD, respectively), as a result of a meta-learning process where the skill of the *Demonstrator* is inferred to modulate imitation.

## Discussion

Over two experiments, we found that whenever imitation is adaptive (*i.e.*, *Skilled Demonstrator*), it takes the computational form of *value shaping*, which implies that the choices of the *Demonstrator* affect the value function of the *Learner*. In other terms, imitation is instantiated as a model-free learning process, as it does not require an explicit model of the *Demonstrator* (15). Our conclusions are based on model comparison and parameter comparison analyses, whose validity is supported by model recovery and parameter recovery quality checks. Our results are robust across experiments and across different implementations of the computational models. In accordance with a vast body of evolutionary literature indicating that, in order to be adaptive, imitation should be modulated (16), we found that humans can correctly infer the skill of a *Demonstrator* and modulate their imitation accordingly (7). Finally, a comparison between reward and imitation learning rates suggests that privately generated outcomes are over-weighted compared to social information.

### Value shaping vs. decision biasing

*Decision biasing* postulates that imitation is essentially a exploration bias rather than a learning process. By contrast, *value shaping* allows the *Demonstrator*’s choices to have a stronger and long lasting influence on the Learner’s behaviour, as demonstrations are progressively accumulated and stored in the value function. A notable advantage of *value shaping* is that it can easily account for observational learning in Pavlovian settings where no decisions are involved; while *decision biasing* cannot (17, 18). Our *value shaping* method is equivalent to the *outcome bonus* method that has been proposed for integrating advice into reinforcement learning (11). This method stipulates that the advised options are perceived more positively, and thus acquire an extra reward bonus. In the reinforcement learning literature, this strategy is called *reward shaping*, which corresponds to augmenting the reward function with extra rewards in order to speed-up the learning process (19). However, it has been widely reported that reward shaping can lead to sub-optimal solutions that fail to account for human behaviour (12, 20, 21). Several solutions have been proposed to address this problem, such as *value shaping* which affects the preference for "advised" actions without modifying the *Learner*’s reward specifications (12), and *policy shaping* which affects the *Learner*’s behaviour without modifying its value function (22, 23)^1^.

In our work, the term *value shaping* can be used interchangeably with *reward shaping* and *policy shaping*. The distinction between *reward shaping*, *value shaping*, and *policy shaping* can not be addressed in single-step problems such as in our task. In fact, adding an extra reward bonus to an action is equivalent to augmenting its expected value, which is equivalent to augmenting its probability of being selected. Further research using multi-step reinforcement learning paradigms is needed to assess which of these three shaping methods best accounts for human behaviour.

### Value shaping vs. model-based imitation

Our results show that human *Learners* imitate a *Skilled Demonstrator* by integrating the observed actions into their own value function, without building an explicit model of the *Demonstrator*. This solution has the obvious advantage of being computationally simpler than *model-based imitation*. In our study, the distinction between *value shaping* and *model-based imitation* echoes the classical distinction between *model-free* and *model-based* methods in the reinforcement learning literature. At first view, our findings may seem in contrast with previously reported results showing that people infer a model of a *Demonstrator* (5, 9) and can successfully predict other’s behaviour (24). These seemingly contradictory findings can be partially explained by the fact that in these works, participants were explicitly instructed to predict other subject’s behaviour; whereas in our tasks, they were not. So it might be that subjects do not build a model of the *Demonstrator* unless there is a need for it, or being explicitly asked for. Moreover, in these previous works demonstrations were the only available source of information for the *Learner*; while in our task, demonstrations were in competition with self-generated outcomes. As the task is defined by the outcomes, they constitute a reference model over which other learning signals might be integrated. *Value shaping* represents a more parsimonious solution than building separate models for every learning signal.

More generally, we can imagine that both learning methods exist and correspond to distinct and co-existing inference modes that are used in different contexts: when only demonstrations are provided, inferring the *Demonstrator*’s goal is the only way for learning the task, whereas in presence of another source of information (*i.e.*, self-generated rewards), demonstrations only play a secondary role.

Which of these learning modes is used for imitation can have distinct implications. *Model-based imitation* implies that people build distinct representations of the same task, one of which can be a representation of a *Demonstrator*’s goal. Each representation can then influence the others, be switched on and off, be more or less considered by the *Learner*. This provides a certain control to the *Learner* on the capacity to disentangle the reliability of each model. *Value shaping*, on the other hand, implies a deeper effect of imitation. People would integrate others’ behaviour and adopt their preferences as their own. This "subversive" effect of imitation can be found in other works (25), where over-imitation is explained not as a mimicry mechanism, but by the modification of the inner representation of the causal model of the environment.

### Adaptive modulation of imitation

Our findings are consistent with the repeated observations in theoretical studies indicating that imitation should be a controlled process (1, 14, 16): for imitation to be adaptive, it should be modulated by several environmental and social signals. Here we focused on the *Demonstrator*’s skills and found that when the *Demonstrator* was not skilled, the imitation rate was downregulated. In other terms, consistent with previous studies, we show that the adaptive regulation of imitation can be achieved by monitoring endogenous signals (likely an estimate of the *Demonstrator*’s performance) and does not necessary require explicit cues and instructions (7, 8, 26). At the computational level, this effect can be formalized as a meta-learning process, where relevant variables are used to optimally tune the learning parameters (27, 28). For example, this could be achieved by tracking the difference between the reward earned following the *Demonstrator*’s choices and opposite choices; then using this relative merit signal to adaptively modulate imitation (29).

Interestingly, subjects in Exp.1 were more successful in modulating their imitation of the *Unskilled Demonstrator* compared to subjects in Exp.2 (cf. Fig. 3c). A possible interpretation could be that subjects in Exp.1 had the opportunity to compare the performance of both *Demonstrators*, and thus avoid imitating the *Unskilled Demonstrator*. In Exp.2, even though subjects imitated the *Skilled Demonstrator* more than the *Unskilled Demonstrator*, the effect was less significant than in Exp.1. This suggests that comparing demonstrations to one’s own performance is not sufficient for spotting bad examples; whereas comparing good and bad demonstrations is important for avoiding bad influence.

Our model parameters analysis also showed that the imitation rate was smaller compared to the reward learning rate, even when the *Demonstrator* outperformed the participant (80% average correct response rate vs. <75%). This finding may suggest that human adults display an egocentric bias, where privately generated outcomes are over-weighted compared to social cues. Whether or not the difference between private and imitation learning rates changes as a function of task related factors (*e.g.*, task difficulty) or inter-individual differences (*e.g.*, age of the participants, pathological states) remains to be established (30–32).

### Putative neural bases

Our computational modeling results suggest that the actions of a *Demonstrator* constitute a pseudo-reward, generating a reward prediction error used for updating the *Learner*’s value function. Previous imaging data provided evidence consistent with this neuro-computational hypothesis. Specifically, Lebreton et al. (33) showed, in a simple option valuation task, that action observation signals (originally encoded in the human mirror neuron system (34)) progressively affect reward-related signals in the ventral striatum and the ventral-prefrontal cortex, two areas robustly associated to reward prediction error encoding in reinforcement learning (35). This neuro-computational hypothesis is also consistent with data showing action prediction errors in the lateral prefrontal cortex (overlapping with the mirror neuron system) and modulation of reward-related signals in the striatum and ventral prefrontal cortex, when following advice (6, 11).

### Conclusions

Our work provides robust evidence that, in the context of social reinforcement learning, imitation takes the form of a computationally simple, yet long lasting, *value shaping* process, where the actions of the *Demonstrator* durably affect the value function of the *Learner*. Imitation plays a key role in the propagation of ideas and behaviours among individuals (36). The global adoption of social media has exacerbated the propagation of different kinds of behaviours, spanning from anodyne memes and lifestyle habits, to more political (37) and economic (38) behaviours. These massive cascade effects can have deleterious impacts, such as political polarization (39), diffusion of fake news (40), and even more macabre trends caused by the propagation of dangerous behaviours (41). Understanding the mechanisms underpinning social influence is therefore crucial for understanding and preventing these negative effects. Our main finding is that imitation can be understood as a modification of *values* rather than a bias over *decisions*. The ensuing persistence of imitation may in part explain the strength and pervasiveness of phenomena related to social influence. Future research will determine whether or not imitation is impaired in social and non-social psychiatric conditions, such as depression, anxiety, autism and borderline personality disorder, and whether these hypothetical impairments take the form of shifts in its computational implementation and/or model parameters (42).

## Methods

### Participants

Respectively 24 and 44 - new - healthy participants were enrolled in our two experiments (Table 1). Participants were recruited through the *Relais d’Information sur les Sciences Cognitives* website^2^. The inclusion criteria were being over 18 years old and reporting no history of psychiatric and neurological illness. All study procedures were consistent with the Declaration of Helsinki (1964, revised 2013) and participants gave their written informed consent, prior to the experiment. Participants were reimbursed 10 to 20 euros for their participation, depending on their performance (on average 13,9 *±* 4.69 euros).

**Table 1.**
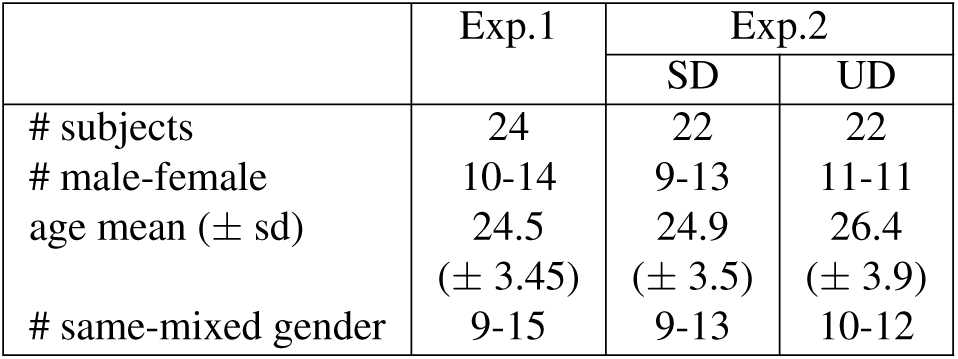
Demographics of the two cohorts of participants.

|  | Exp.1 | Exp.2 |  |
| --- | --- | --- | --- |
|  |  | SD | UD |
| # subjects | 24 | 22 | 22 |
| # male-female | 10-14 | 9-13 | 11-11 |
| age mean (± sd) | 24.5<br>(± 3.45) | 24.9<br>(± 3.5) | 26.4<br>(± 3.9) |
| # same-mixed gender | 9-15 | 9-13 | 10-12 |

### Experimental task and procedure

Both experiments implemented variants of a standard social reinforcement learning task, where, in addition to private learning trials and conditions, the participants were sometimes exposed to the choices of another agent. More details concerning the experimental task are provided in the Result section. Participants were recruited in pairs - either mixed or same gender - and received the instructions jointly. Subjects were tested in two separated cabins, they were asked in advance to provide a electronic version of a passport-like photo (displayed on the upper part of each side of the computer screen), to clearly differentiate private and observational trials. They were told they would engage in a small game in which they could sometimes observe each other’s choice in real time – but not the outcomes associated with that action. Participants were informed that some symbols would result in winning more often than others, and were encouraged to accumulate as much points as possible. It was emphasized that they did not play for competition but for themselves and importantly, there was no explicit directive to observe the other player. The participants were not explicitly instructed to integrate the choices of the other player into their own learning and decision-making, but it was explicit that both a given symbol has the same value for both subjects. At the end of each session, the experimenter came in both cabins to set the subsequent one, and participants were instructed to begin each session at the same time. The debriefing of the experiment was done with each pair of subjects, and all participants were debriefed about the cover story after the experiment.

### Computational modeling

We compared three hypotheses about the computational implementation of imitation in reinforcement learning. All three models are the same for learning from self-experienced outcomes. They only differ in how they integrate demonstrations into the learning process.

#### Baseline model

As a baseline, we implemented a Rescorla-Wagner model that learns only from self-experienced outcomes while ignoring demonstrations. A value function, Q, is first initialized to zero. Then, on every time-step, when the subjects chooses the option *c* and observes an outcome *r*,the value of the chosen option is updated as following:

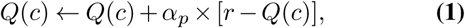

where *α*_*p*_ is a "private" learning rate. Action selection is performed by transforming action values into a probability distribution through a softmax function

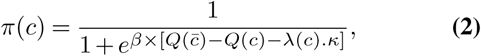

where 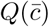 is the value of the unchosen option 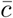, *κ* a choice auto-correlation parameter (43) and *λ*(*c*) is defined as:

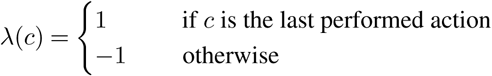

We compared two implementations of this baseline depending on how the Q-values are updated. The first implementation, RW1, uses the *value update* scheme described in Eq. 1. The second implementation, RW2, is based on a *symmetric value update* that also updates the value of the unchosen option in the opposite direction with

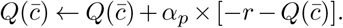

#### Decision biasing

Decision biasing builds on the baseline when it comes to learning from experienced outcomes. However, demonstrations are also used for biasing action selection. We tested six different implementations of this model in order to control for some computational details that may affect the fitting quality (cf. Fig. 4). The first implementation, DB1, is the original model presented in (6) and used in (7). When a demonstration *d* is observed, it is used for biasing the policy *π* as following. First, the policy *π* is derived from the value function *Q* via Eq. 2. Then, the policy *π* is modified via:

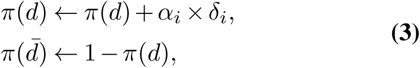

where *π*(*d*) and 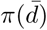 represent respectively the probability to select the demonstrated and the non demonstrated option, and *α*_*i*_ an *imitation rate*.

We refer to this update scheme as *policy update*. In order to control for the fact that in decision biasing the *action prediction error* is computed at the level of the policy and not the Q-values, we devised another version, DB2, in which the action prediction error is computed based on the Q-value (*i.e.*, a *value update*). But still, this update must only affect action selection without modifying the value function. To do so, we keep a *"decision value function"*, *Q′*, which is a copy of the value function *Q* that is used only for action selection. The policy *π* is derived from *Q′* instead of *Q*. When a demonstration *d* is observed, it is used for biasing *Q′* as following. First, *Q′* is copied from *Q*. Then, it is modified via:

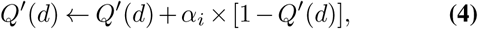

**Fig. 4.**
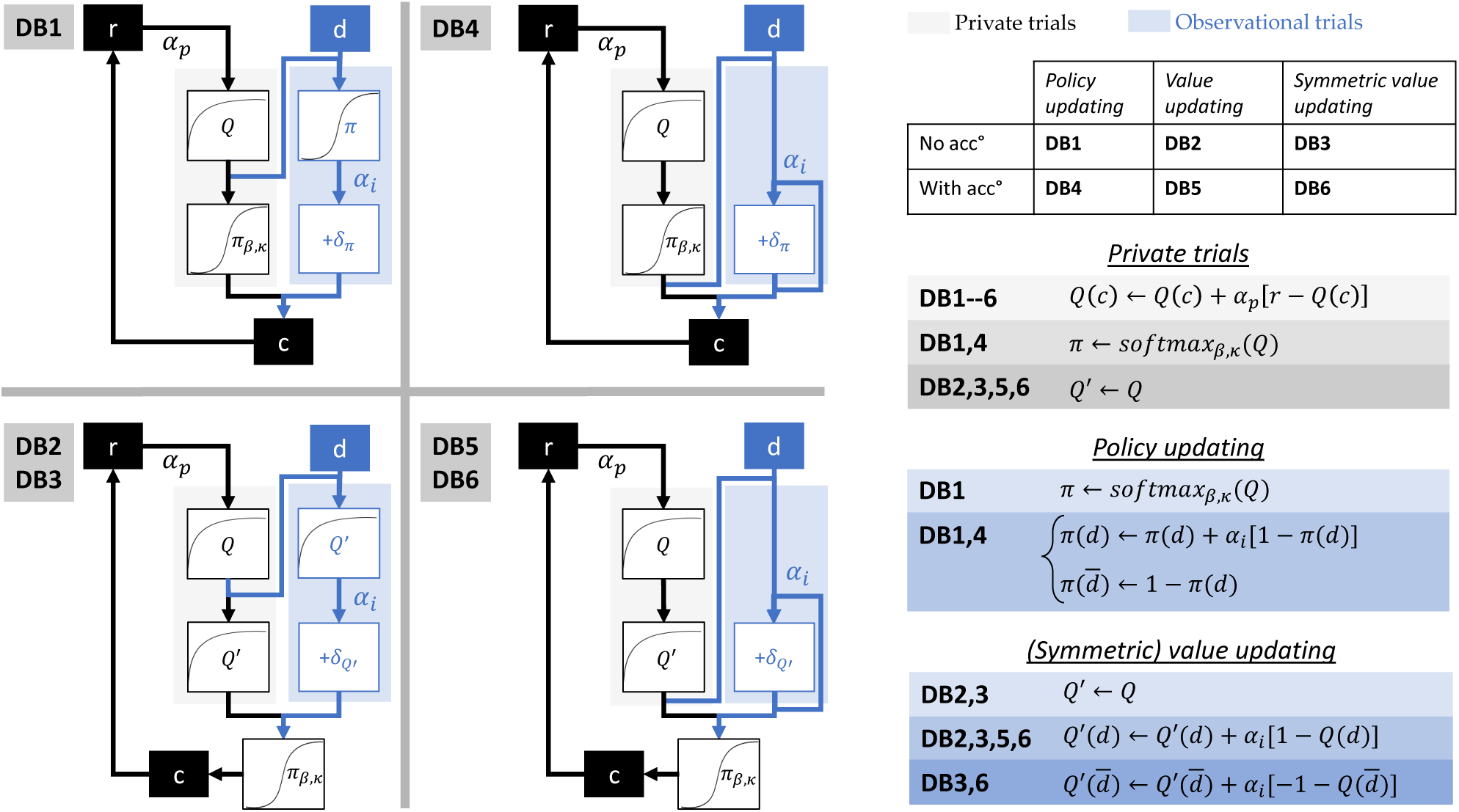
Six implementations of decision biasing. In DB1, demonstrations bias *Learner*’s actions via *policy update*. DB2 and DB3 implement the same mechanism through *value update* and *symmetric value update*. DB4,DB5 and DB6 are equivalent to DB1,DB2 and DB3, while allowing for the accumulation of successive demonstrations. This is done by removing the first step of the update where *π* or *Q′* is derived from *Q*. Accumulation is depicted in the diagrams by the loop within observational trials.

One difference between Eq. 3 and Eq. 4 lies in the symmetry of the update. In Eq. 3, increasing the probability of selecting one option naturally decreases the probability of selecting the other option. However in Eq. 4, increasing the value of one option does not affect the value of the alternative option. To account for this fact, we implemented an extension of DB2, in which we also update the value of the alternative option in the opposite direction. DB3 implements a *"symmetric value update"* via

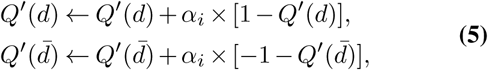

One common limitation of all these three implementations is that they do not allow the accumulation of successive demonstrations, because the policy *π* (resp. *Q′*) is derived from *Q* each time a demonstration is provided. To address this limitation, we extend these implementations, by removing the initial step of the update that derives *π* (resp. *Q′*) from *Q*. This way, successive demonstrations within a same block accumulate their effects.

#### Model-based imitation

Just like *decision biasing*, *model-based imitation* is built on the baseline for learning from self-experienced outcomes. However, demonstrations are used for building a model of the demonstrator’s preferences. This model is then used for biasing action selection. We compared nine different algorithmic implementations of this model (cf. Fig. 5). These implementations differ in how the model of the *Demonstrator* is built and how it is used for biasing action selection. We construct a 3×3 model space based on whether each step uses a *policy update* (Eq. 3), a *value update* (Eq. 4), or a *symmetric value update* (Eq. 5). To note, a *symmetric value update* of the *Demonstrator*’s model corresponds to the *approximate inverse reinforcement learning* algorithm implemented in (9). In all nine implementations, the *Demonstrator*’s model is updated with a learning rate *α*_*d*_, and used for biasing action selection with an *imitation rate α*_*i*_. Finally, for each of these nine implementations, we compared two versions, one where *α*_*d*_ is fitted as a free parameter, and one where it is fixed to 0.1 (cf. supplementary Fig. 5).

**Fig. 5.**
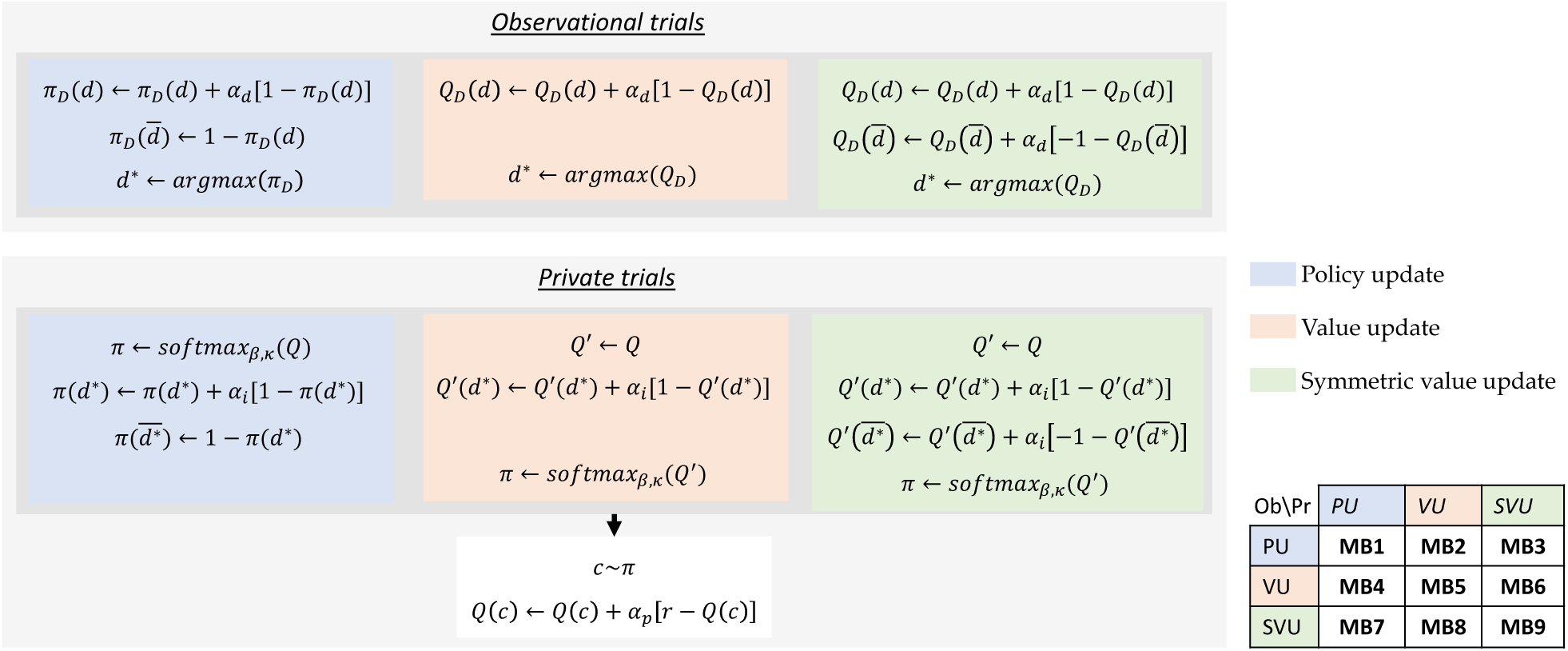
Nine implementations of model-based imitation. In observation trials, demonstrations are used for building a model of the *Demonstrator* (*π*_*D*_ or *Q*_*D*_). This is done either through *policy update* (MB1, MB2 & MB3), *value update* (MB4, MB5 & MB6), or *symmetric value update* (MB7, MB8 & MB9). In private trials, the model of the *Demonstrator* is used for biasing *Learner’s* actions through either *policy update* (MB1, MB4 & MB7), *value update* (MB2, MB5 & MB8), or *symmetric value update* (MB3, MB6 & MB9).

#### Value shaping

In *value shaping*, demonstrations are directly integrated in the *Learner*’s value function by reinforcing the demonstrated actions. We compared two implementations of this model. VS1 performs a *value update* via

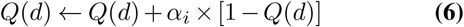

VS2 performs a *symmetric value update* by also updating the value of the non demonstrated option via

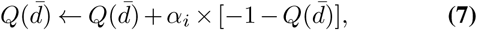

### Model fitting procedure

#### Model comparison

In a first phase, we compared the different implementations of each model. The best implementation was then included in the final model-space for comparison (cf. supplementary Fig. 2).

Each model was fitted separately to the behavioural data of our two experiments. First, the free parameters of each model were optimized as to maximize the likelihood of the experimental data given the model. To do so, we minimize the negative log-likelihood using the *fmincon* function in Matlab, while constraining the free parameters within predefined ranges (0 *< α <* 1, 0 *< β <* + ∞). The negative log-likelihood LL is defined as

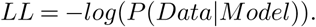

Then, from the negative log-likelihood we derive the Akaike Information Criterion (AIC) (44) defined as

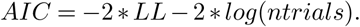

The AIC is -with other metrics-a commonly used metric for estimating the quality of fit of a model while accounting for its complexity. As such, it provides an approximation of the out-of-sample prediction performance (45). We selected the AIC as a metric after comparing its performance in model recovery along with other metrics such as the BIC (see the paragraph about model recovery).

Finally, the individual AIC scores (actually AIC/2) were fed into the mbb-vb-toolbox (46). Contrary to fixed-effect analyses that average the criteria for each model, the random-effect model selection allows the investigation of inter-individual differences and to discard the hypothesis of the pooled evidence to be biased or driven by some individuals – *i.e.* outliers. This procedure estimates the model expected frequencies and the exceedance probability for each model within a set of models, given the data gathered from all participants. The expected frequency is the probability of the model to generate the data obtained from any randomly selected participant – it is a quantification of the posterior probability of the model (PP). It must be compared to chance level, which is one over the number of models in the model space. The exceedance probability (XP), is the probability that a given model fits the data better than all other models in the model space. Theoretically, a model with the highest expected frequency and the highest exceedance probability is considered as ‘the winning model’.

#### Parameter optimization

Models parameters were optimized by maximizing the logarithm of the Laplace approximation to the model evidence (*i.e.*, Log Posterior Probability, LPP)

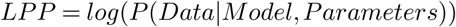

LPP maximization includes priors over the parameters (temperature gamma (1.2,5); LR beta (1.1.,1.1). Essentially, it avoids wrongful fitting of the parameters estimates that could be driven by noise. The same priors were used for all learning rates, to avoid bias in learning rate comparison. Of note, the priors are based on previous literature [70] and have not been chosen for this study.

#### Parameter comparison

We use non parametric tests for comparing model parameters. Wilcoxon signed-rank test is used for comparing paired learning rates, such as private vs. imitation learning rates (Fig.3c) and skilled vs. unskilled imitation learning rates in Exp.1 (Fig.3c). Wilcoxon rank-sum test for non paired learning rates, such as skilled vs. unskilled imitation learning rates in Exp.2 (Fig.3c).

#### Model recovery

The main aim of the model recovery procedure is to verify that the models that we try to compare can effectively be distinguished given the experimental data and the model fitting procedure. It could be the case that two competing models are not sufficiently different from each other to be effectively distinguished, or that the experimental design is not appropriate for eliciting any difference between these models.

Model recovery consists in running several simulations of each model with the historical data of each subject, *i.e.* the observed stimuli and demonstrations. Model parameters are initialized randomly according to their respective prior distributions. This allows us to have a ground truth about which model has generated each piece of data. After simulation, we run the model fitting procedure on the data generated by each model, and report the model frequency and exceedance probability of each competing model. If the exceedance probability of the model that has generated the data passes a predefined threshold, it means that our model fitting procedure allows us to recover the true generating model.

#### Parameter recovery

Parameter recovery is a useful way to assess the quality of the parameter fitting procedure. To do this, we computed the Spearman correlation between parameter values that were used for generating simulated data, with the values recovered by the model fitting procedure. A high correlation score indicates a reliable parameter fitting procedure.

## ACKNOWLEDGEMENTS

SP is supported by an ATIP-Avenir grant (R16069JS) Collaborative Research in Computational Neuroscience ANR-NSF grant (ANR-16-NEUC-0004), the Programme Emergence(s) de la Ville de Paris, the Fyssen foundation and the Fondation Schlumberger pour l’Education et la Recherche (FSER)

## Supplementary material

**Fig. 1.**
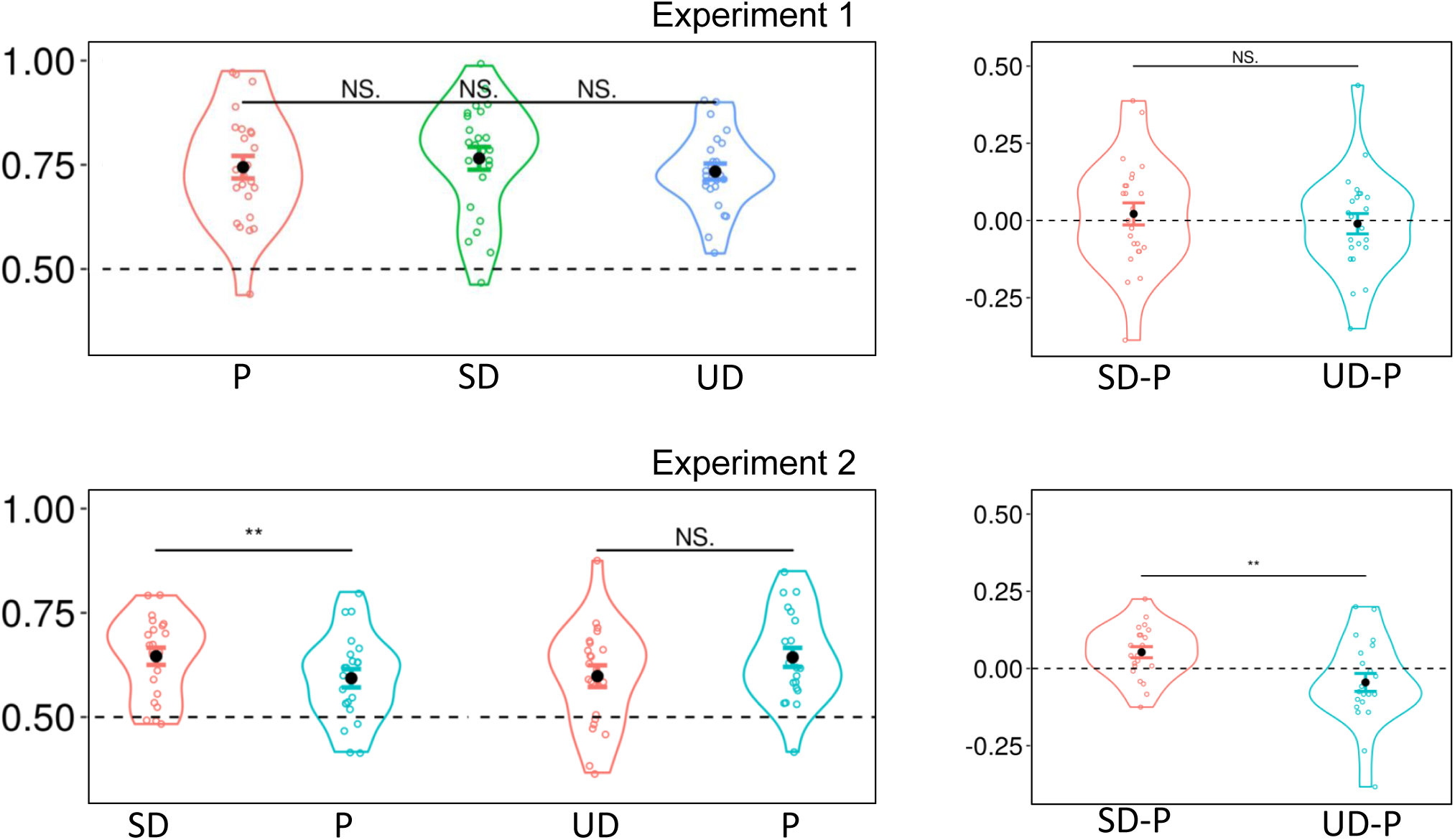
Raw task performance on both experiments, with respect to conditions. P private condition. SD and US observational conditions, for *Skilled Demonstrator* and *UnSkilled Demonstrator* respectively. In Exp.1, in the condition (SD) was higher than in the Private condition (P), which itself was higher than performance in the *UnSkilled Demonstrator* condition (SD). However, this difference in performance was not statistically significant. In Exp.2, the differential in task performance between observational and private conditions was statistically different between the SD and the UD group.

**Fig. 2.**
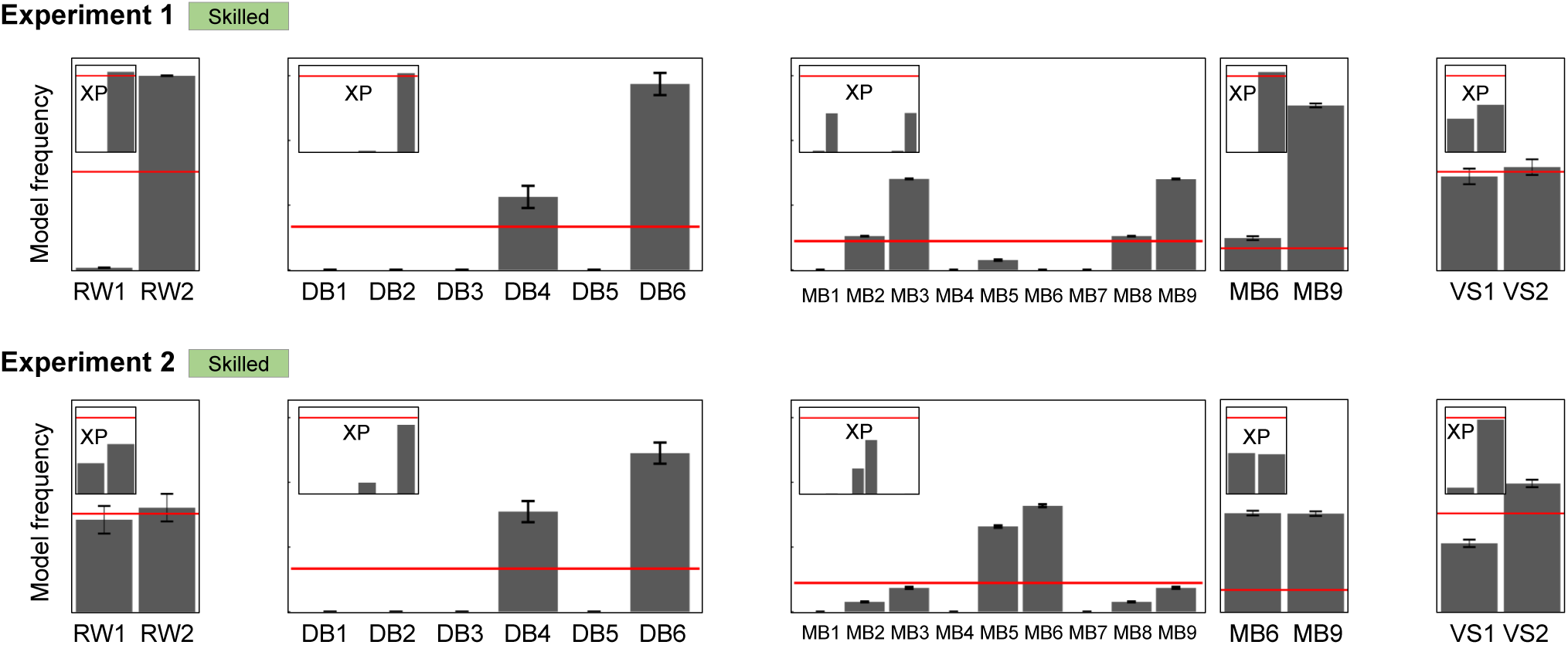
Intermediate model comparisons. We compared different implementations of each model: two for RW, six for DB, nine for MB, and two for VS. For the baseline, implementing a symmetric value update (RW2) improves the model fitting quality in Exp.1. For *decision biasing*, the the best fitting performance in achieved when allowing for symmetric value update from observed demonstrations and for the accumulation of successive demonstrations (DB6). For *model-based imitation*, the best implementations when use the model of the *Demonstrator* for biasing *Learner*’s actions through symmetric value update (MB3 and MB9 in Exp.1, and MB6 in Exp.2). A finer comparison between MB3 and MB9 in both experiments shows that these models are equivalent (not shown here). A further comparison between MB6 and MB9 in both experiments shows that MB9 fits better than MB6. Finally, the best *value shaping* implementation, VS2, uses symmetric value update from observed demonstrations. As a result of this analysis, the final model space includes RW2, DB6, MB9 and VS2.

**Fig. 3.**
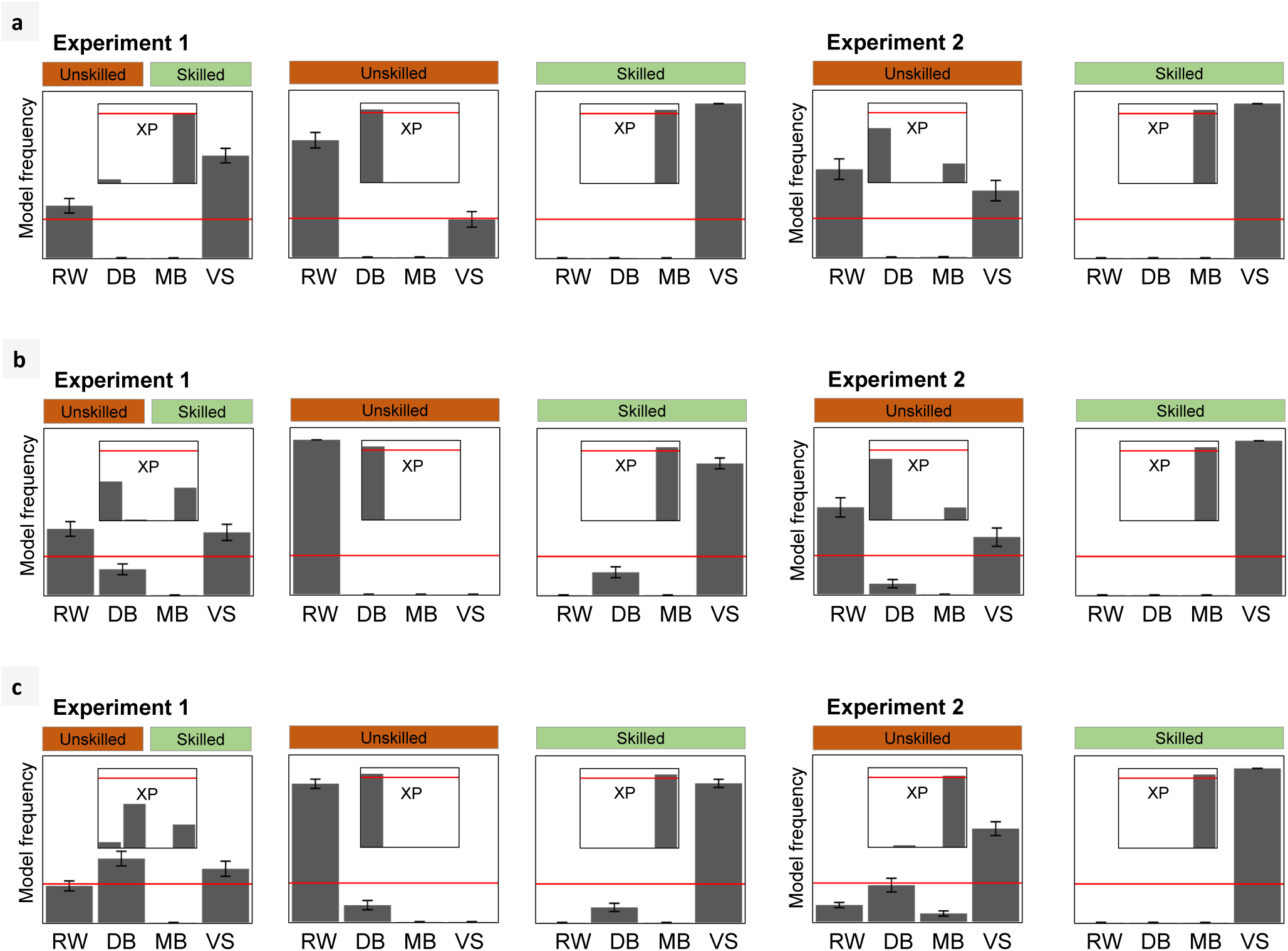
Model comparison results are robust amongst different implementations of the model space. (a) Model space implementation without choice auto-correlation parameter. (b) Model space implementation without symmetric value update for private learning. (c) Model space implementation allowing for negative imitation learning rates. Note that in Exp.2, when allowing for negative learning rates, the winning model in the UD condition is no longer RW, but VS.

**Fig. 4.**
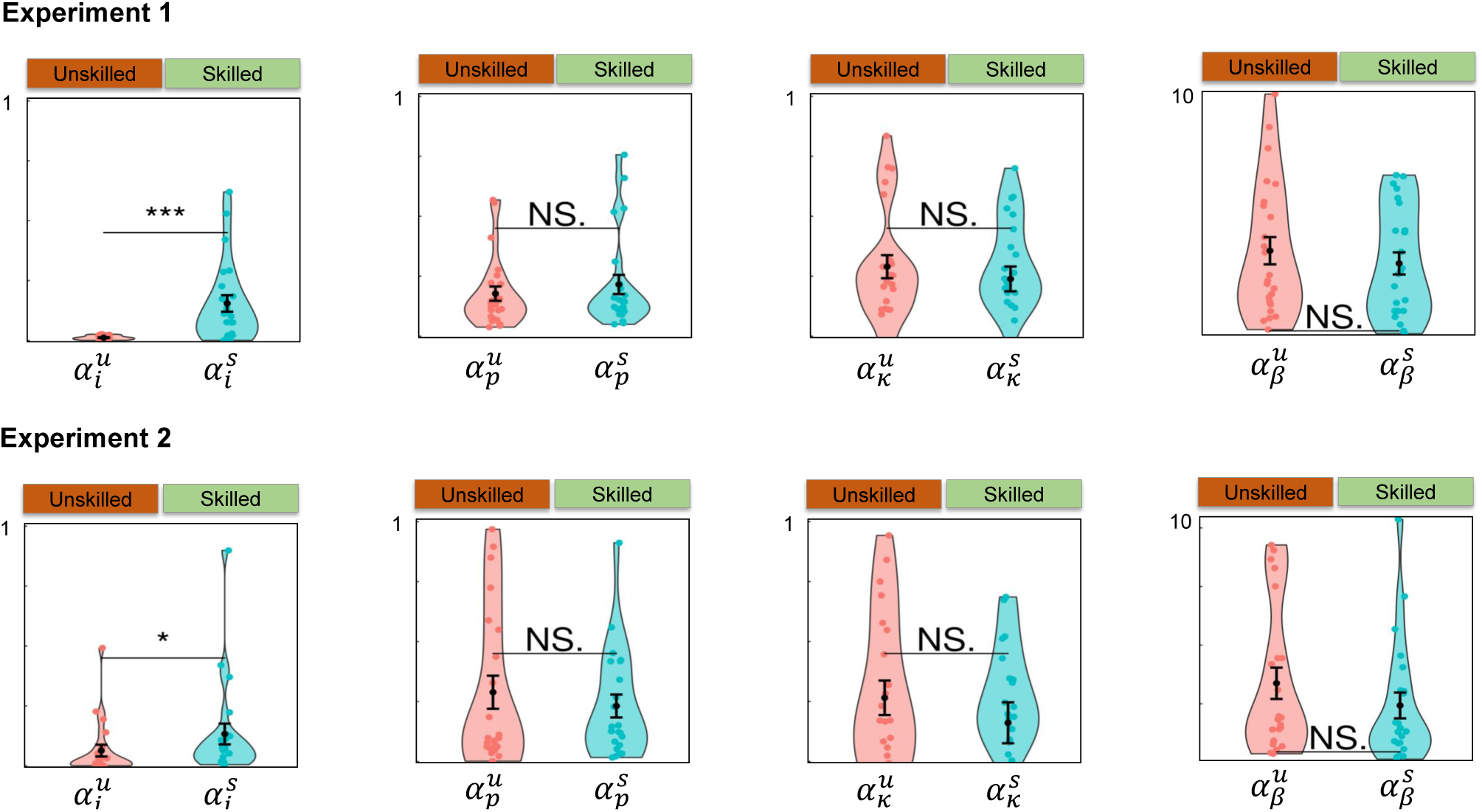
Parameter comparison between UD and SD conditions. No statistical difference was found for non-social parameters *α*_*p*_, *κ* and *β*. Only the imitation learning rate *α*_*i*_ was statistically different across observational conditions in both experiments.

**Fig. 5.**
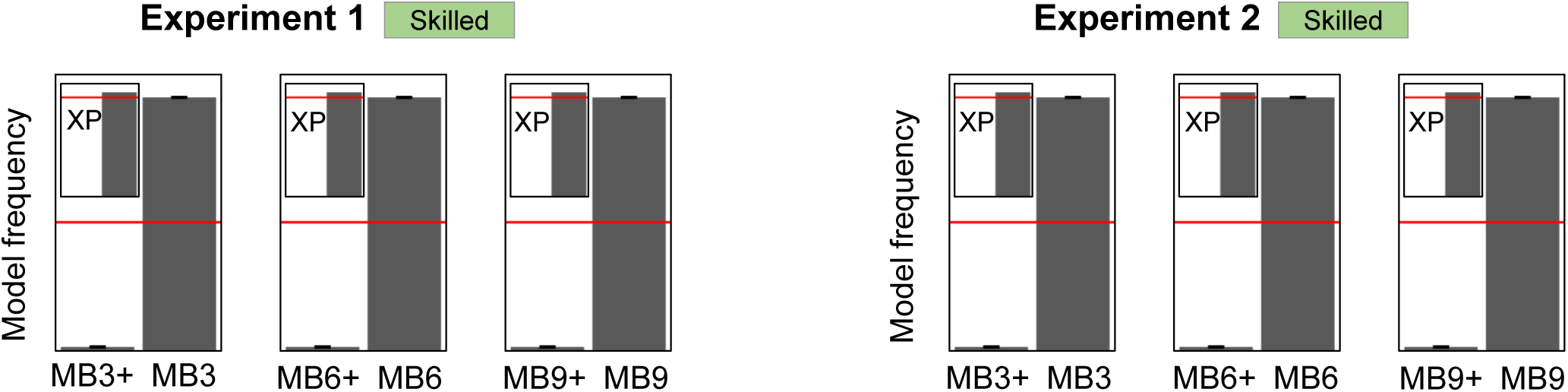
Model comparison between alternative implementations of *model-based imitation* (MB). Specifically, implementations with a fixed parameter *α*_*d*_ = 0.1 (MB3,MB6 and MB9) are better than implementations with *α*_*d*_ as a free parameter (MB3+,MB6+ and MB9+, respectively). *α*_*d*_ is the learning rate used for inferring the preferences of the *Demonstrator*.

## Footnotes

1 Note that *policy shaping* is different from *decision biasing* as it has a long-lasting effect.

2 https://www.risc.cnrs.fr/

